# Enhancement of spatial learning by 40 Hz visual stimulation requires parvalbumin interneuron-dependent hippocampal neurogenesis

**DOI:** 10.1101/2024.04.28.591481

**Authors:** Hai Yan, Xufan Deng, Yunxuan Wang, Shiyu Wu, Jun Du, Mei Yu, Bo Liu, Huimei Wang, Yifan Pan, Zhengyu Zhang, Jinghong Chen, Yizheng Wang, Tara Walker, Perry Bartlett, Jun Ju, Sheng-Tao Hou

## Abstract

Acute and short-term rhythmic 40 Hz light flicker stimulation has shown promising results in alleviating cognitive impairments in mouse models of Alzheimer’s disease (AD), stroke, and autism spectrum disorders (ASD). Understanding the long-term impacts and underlying mechanisms is crucial to progress this approach for potential human therapeutic applications. Here, we show that prolonged exposure to 40 Hz light flicker (1 hour per day for 30 days) significantly improved spatial learning and neurogenesis in the dentate gyrus (DG) without harmful behavioral side effects. Mice with transgenic deletion of doublecortin-positive cells (DCX^DTR^) in the adult hippocampus failed to exhibit enhanced neurogenesis and spatial learning with 40 Hz stimulation. Inactivation or knockout of GABAergic parvalbumin (PV) interneurons reduced the effects of 40 Hz entrainment and neurogenesis enhancement. Mechanistically, the stimulation did not alter the regional microvessel blood flow but significantly raised PV excitability and GABA levels and enhanced inhibitory transmission in the DG. Blocking GABA_A_ receptors reversed the improvements in spatial learning and neurogenesis. These data showed that long-term exposure to 40 Hz light flicker enhances spatial learning through PV-dependent adult neurogenesis, which requires elevated GABA as a critical neurochemical mechanism for sustaining adult neurogenesis.

**In brief:** Rhythmic 40 Hz light flicker alleviates cognitive impairments in diverse animal models of neurological diseases. Understanding its long-term effects and mechanisms is vital for advancing its therapeutic potential for humans. Here, we show that prolonged exposure to this flicker improves spatial learning and boosts adult neurogenesis in the dentate gyrus. Activation of PV interneurons and GABAergic support for the newborn immature neurons underlie this effect, demonstrating lasting benefits for the treatment of neurological diseases.

**Highlights:**

- Long-term exposure to 40 Hz light flicker significantly improved spatial learning and neurogenesis in the hippocampus, devoid of adverse effects.
- The 40 Hz light flicker evoked adult neurogenesis requires the activity of GABAergic inhibitory PV interneurons.
- The 40 Hz light flicker raised GABA levels and enhanced inhibitory transmission in the DG.
- Increased GABA serves as a vital neurochemical mechanism to support adult neurogenesis.

## INTRODUCTION

Recent studies of our own and others have demonstrated significant beneficial effects of rhythmic 40 Hz light flicker in alleviating cognitive deficits in animal models of AD^1–3^, stroke^4^, and ASD^5^. While the specific underlying mechanisms vary and are currently under rigorous investigation and debate^6^, several potential pathways have emerged, including the enhancement of mechanisms involved in the clearance of toxic materials^1–3^, improvement in synaptic connectivity and strength^4^, and increased neurochemical mediators^5,7^. Our human brain electroencephalogram studies have confirmed not only an increase in 40 Hz entrainment in various brain regions but also a substantial alteration in microstates associated with brain functions damaged in AD and stroke patients^8^. Importantly, the 40 Hz light flicker has received expedited approval from the FDA and shown promising results in Phase II clinical trials involving AD patients^9^. Despite this progress, one crucial consideration in the abovementioned studies is that the light flicker treatment was acute and short-term, typically lasting less than 14 days. To advance this remarkably straightforward treatment for human therapy, it is crucial to thoroughly understand the long-term effects and mechanisms of action. To this end, we systematically treated adult mice with long-term 40 Hz light flicker (1 hour per day for 30 days) and examined the impact and mechanism affecting the altered cognitive functions.

In the adult mouse brain, neurogenesis predominantly occurs in the hippocampal DG region and the subventricular zone (SVZ) of the lateral ventricles. Generating new neurons in the DG region during adulthood is crucial for learning and memory formation^10–13^. Newly formed neurons in the DG, identified by the expression of the DCX protein, migrate from the stem cell niche in the subgranular layer to the granular cell layer. After a limited number of divisions, they integrate into the existing neural circuits. However, neurogenesis in the DG region diminishes with age in adulthood^14,15^. In contrast, newborn neurons originating in the SVZ migrate along the outer wall of the lateral ventricles toward the olfactory bulb. There, they develop into olfactory inhibitory interneurons and become part of the neural networks^16^.

In the DG, specific markers distinguish different stages of neurogenesis. For instance, the expression of SOX2 characterizes non-radial and horizontal type-1 cells. T-brain 2 (TBR2) is expressed in type-2a and type-2b intermediate neuroprogenitor cells that eventually mature into neuroblasts (type-3)^17^. DCX is an early marker expressed in immature granule cells during migration and dendritic growth, but its expression is deactivated before these neurons reach maturity^18–21^. These immature neurons then establish synaptic connections with existing mature neurons and gradually integrate into the neural circuit^22,23^.

The precise regulation of neurogenesis in the hippocampus is heavily reliant on localized neural circuits that integrate individual experiences, neural activity, and regulatory mechanisms of neurogenesis^24^. Within these circuits, two primary types of GABAergic interneurons play a vital role in supporting the development of newborn neurons. These are the fast-spiking PV interneurons and somatostatin-expressing SST interneurons, both of which inhibit the dendrites of granule cells. PV interneurons are typically situated closer to the granule cell layer and provide lateral and recurrent inhibition, facilitating continuous GABA release^25^. Synaptic connections between PV interneurons and newborn neurons can be observed as early as 7 days postnatally, preceding the connections established by SST interneurons^26^. Previous studies have shown that optogenetic inhibition of PV interneurons, but not SST interneurons, effectively suppresses neurogenesis in the DG region^27^.

Activation of PV interneurons may function as sensors of circuit activity and provide local cues to facilitate the various components within the hippocampal neurogenic niche. Of particular note is the impact of GABA, which, through spillover effects and subsequent synaptic release to immature neurons, effectively elevates the resting potential of newborn neurons, making them more responsive to action potentials^28–31^. This excitatory effect of the inhibitory neurotransmitter GABA on newborn neurons is attributed to the abundant expression of NKCC1 chloride channels, resulting in a depolarized equilibrium potential for chloride ions in newborn neurons, particularly within the initial three weeks postnatally^26,32,33^.

Here, we discovered that long-term 40 Hz light flicker entrainment promoted DG neurogenesis and enhanced spatial learning. Enhancement of GABA release through the activation of PV interneurons is required for the inhibitory transmission to sustain the maturation of DCX^+^ cells in the adult mouse DG.

## RESULTS

### Long-term 40 Hz light flicker stimulation significantly enhanced spatial learning

To assess the impact of 40 Hz light flicker treatment on cognitive functions, 6-month-old mice underwent daily exposure to 40 Hz light flicker for 1 hour for 30 days. Same-age control mice were maintained under the same light-and-dark (12:12 hours) lighting cycle conditions without such intervention (Figure 1A).

**Figure 1.**
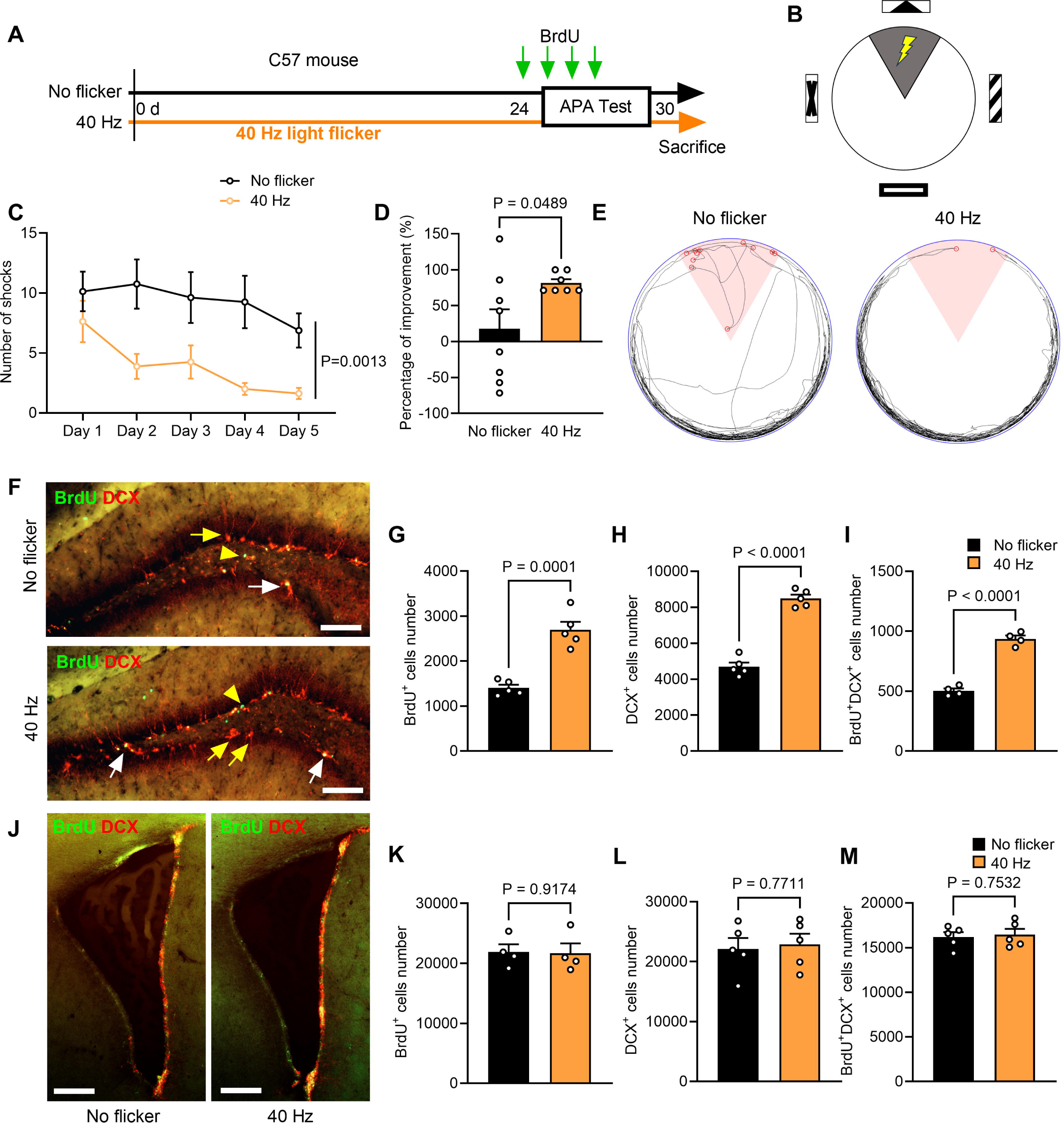
Long-term 40 Hz light flicker treatment significantly enhanced DG spatial learning and neurogenesis. (A) The experimental design and timeline schema show “No flicker” and “40 Hz” groups. (B) The schematic diagram of the APA test. (C) The long-term 40 Hz light flicker-treated mice (40 Hz group) received significantly fewer shocks than the No flicker group of control mice during the APA test (n = 8 mice for no flicker group and n = 7 for the 40 Hz group). A two-way RM ANOVA with Tukey’s *post hoc* test was performed to analyze the effect of 40 Hz light flicker treatment and the days on the number of shocks. The result revealed a statistically significant interaction between the effects of 40 Hz stimulation and the number of shockers received [F(14, 1) = 15.96, P = 0.0013]. (D) The percentage improvement in avoiding shocks on the 5^th^ day of APA testing compared to the 1^st^ day of APA test. (E) The representative movement trajectory map on the 5^th^ day of the APA test. The 60° pink area indicates the shock zone/ The red circles indicate where electric shocks were delivered. (F, J) Double immunostaining for BrdU (green) and DCX (red) of the DG (F) and the SVZ (J) of different groups of mice. The yellow-colored arrows indicate DCX^+^ immature neurons born before BrdU labeling. The white-colored arrows indicate BrdU^+^/DCX^+^ cells and the yellow-colored arrowheads show BrdU^+^ cells that are not yet immature neurons. Scale bars, 100 μm. Long-term 40 Hz light flicker treatment significantly increased the number of BrdU^+^ cells, DCX^+^ cells, and BrdU^+^/DCX^+^ cells in the DG (G – I) but not in the SVZ (K – M). Student’s t test was performed for panels D, G-I, and K-M with specific P values as indicated.

The spatial learning performance of these mice was measured using the active place avoidance (APA) test from day 25 to day 30 (Figure 1A). In this task, mice were placed on a rotating platform and evaluated based on their ability to avoid a stationary shock zone using spatial visual cues (Figure 1B). Mice subjected to long-term 40 Hz light flicker treatment showed significant improvement in avoiding the shock zone compared to the untreated mice from test day 2 to day 5 (Figures 1C–1E). Furthermore, the prolonged exposure to 40 Hz light flicker neither induced anxiety-like behavior nor modified the spontaneous motor activity of these mice (Figures S1A-S1C).

### Long-term 40 Hz light flicker treatment significantly enhanced DG neurogenesis

Given the established link between enhanced spatial learning and increased hippocampal neurogenesis^12,34,35^, we designed experiments to determine whether prolonged exposure to 40 Hz light flicker promotes neurogenesis in the adult brain. Mice were injected with BrdU injection daily from day 24 to day 27, following which the number of newly generated neurons in the hippocampal DG and SVZ was evaluated through immunostaining for BrdU and the immature neuronal marker DCX (Figure 1A). In comparison to the untreated mice, long-term 40 Hz light flicker-treated mice exhibited a significant increase in the number of cells positive for staining with BrdU^+^, DCX^+^, and BrdU^+^/DCX^+^ co-staining in the DG region (Figures 1F-1I), while no changes occurred in the SVZ region (Figures 1J-1M). These data suggest a potential linkage between DG neurogenesis and enhancement of spatial learning following long-term exposure to 40 Hz light flicker treatment.

### The enhanced spatial learning conferred by 40 Hz light flicker treatment is neurogenesis-dependent

To investigate whether the enhanced spatial learning by long-term 40 Hz light flicker treatment required adult neurogenesis, we utilized transgenic knock-in DCX^DTR^ mice expressing the human diphtheria toxin receptor (DTR) under the control of the DCX promoter. The deletion of newborn neurons was accomplished by administering diphtheria toxin (DT). DCX^DTR^ mice were subjected to intraperitoneal (i.p.) injections of saline or DT once every 2 days for 2 weeks starting from day 8 (Figure 2A). The open field test (OFT) demonstrated that DT and 40 Hz light flicker did not elicit anxiety-like behavior or affect the spontaneous motor activity of DCX^DTR^ mice (Figures S1D and S1E). Interestingly, the DCX^DTR^+40 Hz group significantly increased their ability to avoid the shock zone during the APA test compared to the DCX^DTR^ group. In contrast, there was no difference in the number of shocks received between the DCX^DTR^ + DT+ 40 Hz group and the DCX^DTR^ + DT group, indicating a lack of learning to avoid the shock zone (Figures 2C and 2D). Moreover, DT-induced deletion of DCX^+^ immature neuron significantly impaired spatial learning (Figures 2B-2D). These findings demonstrated that DCX-targeted elimination of neurogenesis in adult mice abolishes the spatial learning enhancement induced by 40 Hz light flicker treatment.

**Figure 2.**
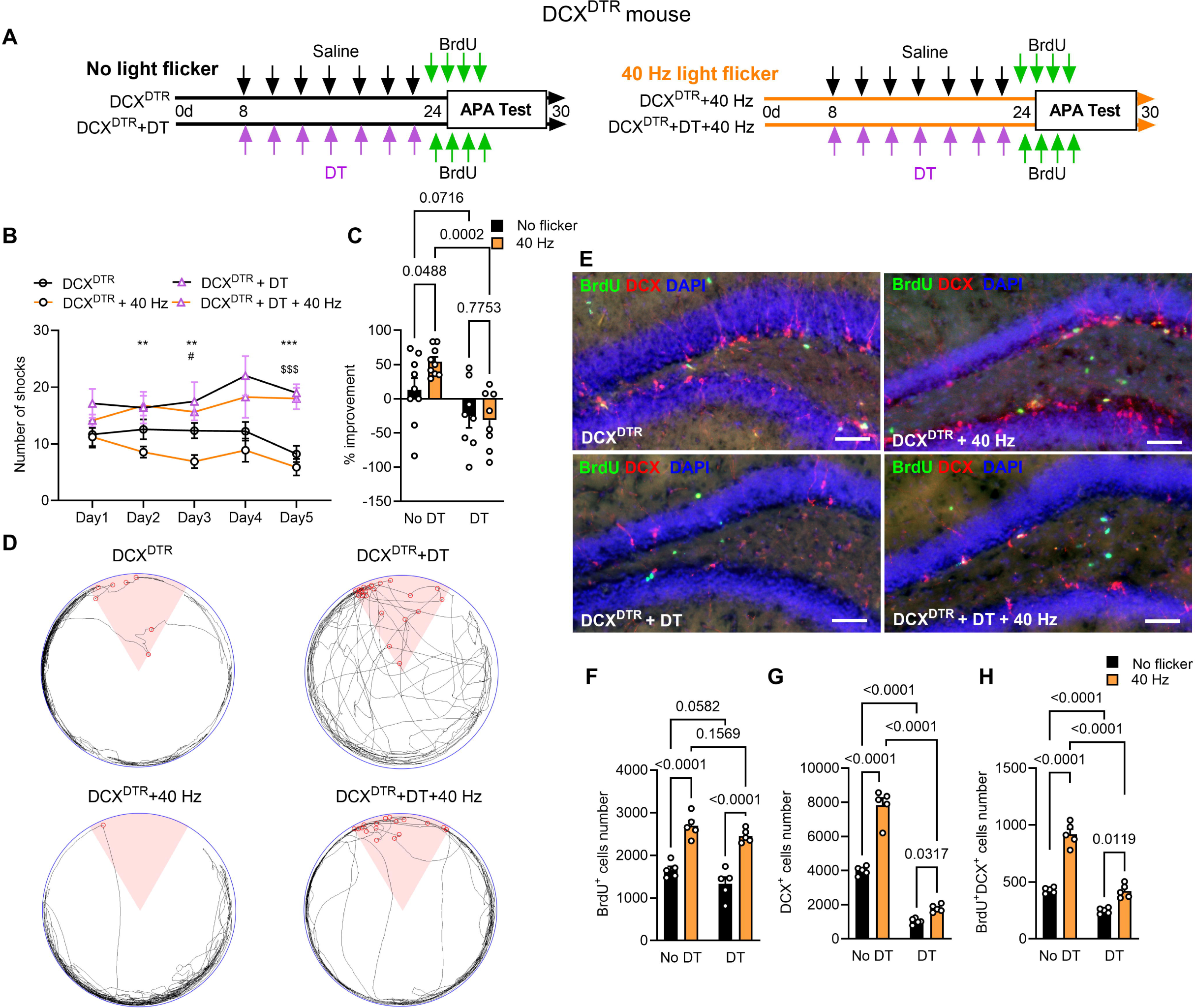
Ablation of DCX^+^ cells in adult mice eliminates 40 Hz light flicker conferred enhancement of spatial learning. (A) Experimental design and timeline schema showing four groups of DCX^DTR^ mice receiving saline (DCX^DTR^), DT (DCX^DTR^+DT), and 40 Hz light flicker (DCX^DTR^+40 Hz and DCX^DTR^+DT+40 Hz). (B) The number of shocks received by the four groups of mice during the last 5-day period of the APA test to determine enhanced spatial learning. (C) The percentage improvement in avoiding shocks on the 5^th^ day of APA testing compared to the 1^st^ day of APA test. DT-induced deletion of DCX^+^ cells in the adult DCX^DTR^ mice significantly eliminated the improvement in spatial learning after long-term 40 Hz light flicker treatment. (D) The representative movement trajectory map on the 5^th^ day of the APA test for the four groups of mice. (E) Double immunostaining for BrdU (green) and DCX (red) with DAPI (blue) of the DG region of the four groups of mice. Scale bars, 100 μm. (F-H) DCX^DTR^ mice treated with DT significantly decreased the number of BrdU^+^ (F), DCX^+^ (G), and BrdU^+^/DCX^+^ (H) cells in the DG. n = 9 mice in the No DT groups and n = 8 mice in the DT groups. Two-way ANOVA with Tukey’s *post hoc* test was used for panels B, C, and F-H with P values as indicated on the graph. For panel B, the specific symbols indicate selective comparisons: *P_DCX_^DTR^_+40Hz vs. DCX_^DTR^_+DT+40Hz_; #P_DCX_^DTR^ vs. _DCX_^DTR^_+40Hz_; $P_DCX_^DTR^ vs. _DCX_^DTR^_+DT_; **P < 0.01, ***P < 0.001; #P < 0.05; $$$P < 0.001.

To confirm the effect of deleting DCX cells on hippocampal neurogenesis in response to light flicker, we evaluated the number of BrdU^+^, DCX^+^, and BrdU^+^/DCX^+^ cells using immunohistochemistry (Figure 2E). The DCX^DTR^ + 40 Hz group exhibited a significant increase in BrdU^+^, DCX^+^, and BrdU^+^/DCX^+^ cells in the DG region compared to the DCX^DTR^ group (Figures 2E and 2F-H). There was no substantial variance in the number of BrdU^+^ cells in the DG between the DCX^DTR^ + DT group and the DCX^DTR^ group, indicating that deletion of DCX in the adult mouse DG did not affect neuroprogenitor proliferation (Figure 2F). In contrast, the number of DCX^+^ cells and BrdU^+^/DCX^+^ cells of the DCX^DTR^ + DT group were significantly lower than those in the DCX^DTR^ group, confirming the efficacy of the DT-mediated DCX^+^ cell depletion (Figures 2G and 2H). Moreover, the DCX^DTR^ + DT + 40 Hz group exhibited no significant alteration in the number of BrdU^+^ cells but displayed a significant reduction in the number of DCX^+^ cells and BrdU^+^/DCX^+^ cells in the DG compared to the DCX^DTR^ + 40 Hz group (Figures 2F-2H). Therefore, the deletion of DCX^+^ cells induced by DT treatment abolished the enhancement of neurogenesis caused by 40 Hz light flicker treatment.

### Long-term 40 Hz light flicker treatment does not alter regional blood flow

To understand the possible mechanisms of long-term 40 Hz stimulation, the DG regional microvessel blood flow was examined using an implanted GRIN lens and two-photon microscopy (Figures S2A-S2D). We found that the blood vessel lumen diameter and the red blood cell velocity of arteriole and capillary in the DG did not change (labeled as 40 Hz) compared with the untreated control (labeled as control) after long-term 40 Hz light flicker treatment (Figures S2E-S2J). The sensitivity of microvessels to light flicker also remained unchanged, as shown by measurement of lumen diameter and red blood cell velocity of arteriole and capillary during flashing of 40 Hz light flicker to the long-term 40 Hz-treated mice (labeled as 40Hz+flickering) (Figures S2E-S2J). These data indicated that long-term 40 Hz light flicker did not affect microcirculation in the DG.

### Activation of DG PV interneurons after 40 Hz light flicker treatment

Considering the crucial function of PV interneurons in promoting neurogenesis^27,36^, we investigated whether the 40 Hz light flicker treatment activated PV interneurons in the DG. Transgenic PV^Cre^ mice received stereotactic injections of rAAV-DIO-jGCaMP7f virus in the DG region. The GCaMP7f fluorescence signals of PV interneurons were captured using fibre photometry. Comparing the fluorescence intensity of PV interneurons before and after the 40 Hz light flicker treatment, we observed a significant increase in GCaMP7f fluorescence intensity (Figures 3A and 3B). This *in vivo* fibre photometry data unequivocally demonstrates the activation of PV interneurons by 40 Hz light flicker treatment.

**Figure 3.**
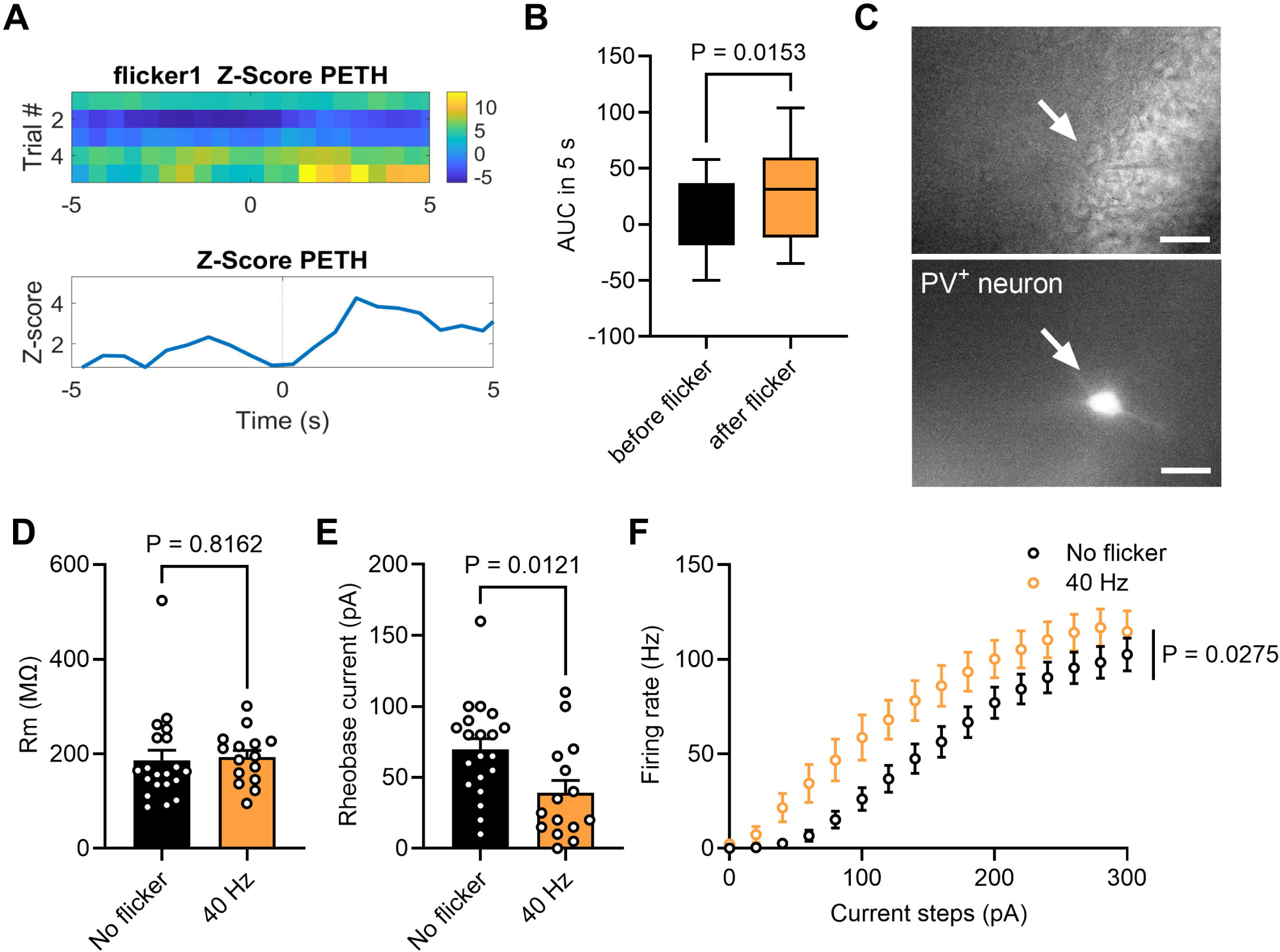
Activation of PV interneurons in the DG region after 40 Hz flicker treatment. (A) Calcium signals of PV interneurons were recorded 5 seconds before and after the 40 Hz light flicker treatment using fiber photometry. The top panel represents the peri-event time histogram (PETH) heatmap of robust z-score signals of individual trials, and the bottom panel represents the averaged z-score signals of PV interneurons of all trials. (B) The fluorescence intensity of PV interneurons significantly increased 5 seconds after the 40 Hz light flicker treatment compared to the 5 seconds before. (C) Patch clamp recording of a PV interneuron viewed using the microscope under phase contrast bright field (top panel; arrow indicates the glass recording pipette tip and the cell) and the epifluorescence field (bottom panel; arrow indicates the PV interneuron). Scale bars, 20 μm. (D, E) Long-term 40 Hz light flicker treatment did not change the R_m_ of PV interneurons (D); however, the rheobase currents decreased (E). (F) The firing rate increased in PV interneurons after the long-term 40 Hz flicker treatment. The Students’ t test was used on panels B, D, and E. A two-way RM ANOVA with Tukey’s *post hoc* test was performed for panel F to analyze the effect of the 40 Hz light flicker treatment and the current on the firing rate. There was a statistically significant interaction between the effects of 40 Hz stimulation and firing rate [F(495, 15) = 1.97, P = 0.0275]. Specific P values were indicated in the panels.

Next, we examined whether prolonged exposure to 40 Hz light flicker treatment affects the electrophysiological properties of PV interneurons. The extended 40 Hz light flicker did not induce changes in the membrane resistance of PV interneurons (Figure 3D). However, a decrease in rheobase currents and an increase in firing rate occurred in these interneurons (Figures 3E-3F). These findings suggest that the extended 40 Hz light flicker treatment influenced the excitability of PV interneurons by lowering their threshold for activation and enhancing their firing capability.

### Deletion and inactivation of PV interneurons reduce gamma entrainment and neurogenesis

To investigate the role of PV interneurons in mediating the effect of 40 Hz light flicker treatment, PV interneurons were specifically deleted using PV^Cre^ mice injected with the rAAV2/9-CAG-DIO-taCaspase3-TEVp-WPRE-pA virus (PV-Cre+Casp3 group). The PV^Cre^ wildtype littermates (without Cre recombinase expression) were injected with the same rAAV (C57+Casp3 group) as a comparative control group (Figure 4A). Local field potential (LFP) was recorded in the molecular layer of the DG (Figures 4B and 4I). The 40 Hz light flicker treatment evoked LFP entrainment in the DG of the C57+Casp3 group and the PV-Cre+Casp3 group, as shown by increased power spectrum density (PSD) at 40 Hz (Figure 4B). However, the PV interneuron deletion group of mice (PV-Cre+Casp3 group) showed a significant decrease in the PSD at 40 Hz frequency (Figure 4C) and the low and high gamma frequencies (Figures 4G and 4H). The theta, alpha, and beta frequency oscillations did not change in the PV-Cre+Casp3 group compared to the C57+Casp3 group following the 40 Hz light flicker (Figures 4D-4F). This data showed that PV interneurons played an important role in mediating the 40 Hz light flicker entrainment in the DG.

**Figure 4.**
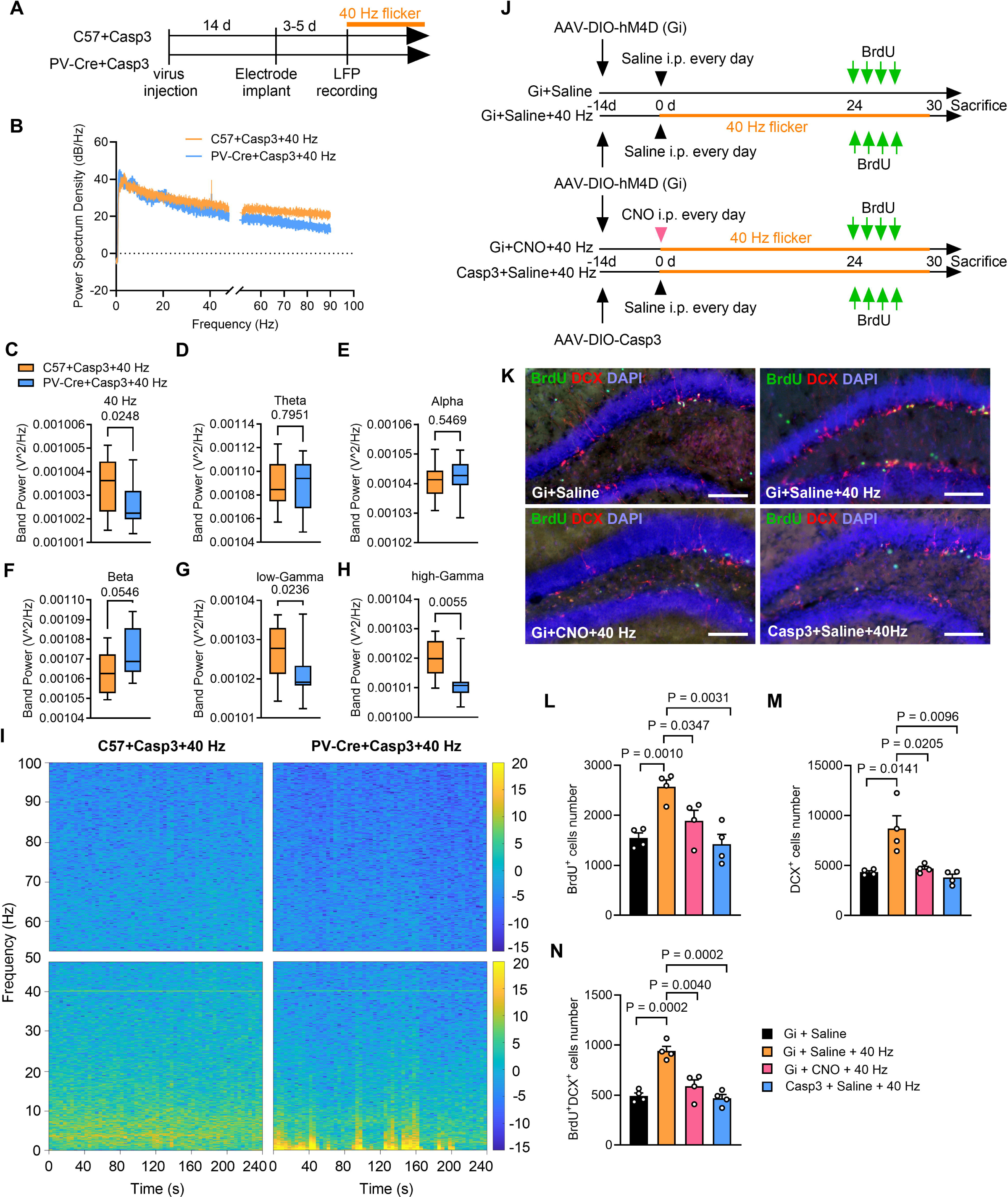
Inactivation of PV interneurons in the DG region reduces 40 Hz entrainment and neurogenesis. (A) An experimental timeline showing deletion of PV interneurons and determination of LFP changes. (B) The PSD of LFP was recorded in the molecular layer of the DG from control mice (C57+Casp3+40 Hz) and PV knockout mice (PV-Cre+Casp3+40 Hz) during 40 Hz light flicker. (C-H) The band power of 40 Hz and the low and high gamma in PV knockout mice were significantly lower than those of the control mice evoked by 40 Hz light flicker. (I) The short-time Fourier transform (STFT) heatmap was generated to show LFP power recorded at 240 s in the DG during the resting state of two groups of mice. The PSD intensity was color-coded according to the scale bars on the right. (J) Experimental schema for inactivation of PV interneurons using inhibitory virus [AAV-DIO-hM4D(Gi) group] and knockout PV interneurons using chemogenetic activation of caspase-3 (AAV-DIO-Casp3 group). (K) Double immunostaining for BrdU (green), DCX (red) with DAPI (blue) of the DG of four groups of mice. Scale bars, 100 μm. (I-N) Inactivation and knockout PV interneurons in PV^Cre^ mice significantly decreased the number of BrdU^+^, DCX^+^, and BrdU^+^/DCX^+^ cells in the DG. Student’s t test was used for panels C-H and L-N. Specific P values were indicated in the panels.

Next, we investigated whether PV interneuron promotes DG neurogenesis after prolonged 40 Hz stimulation using two approaches; one is to use the above-described caspase-3 rAAV to delete PV (Casp3+Saline+40 Hz group), and the second is to use chemogenetic inactivation of PV interneurons (Gi+CNO+40 Hz group). To do this, PV^Cre^ mice received stereotactic injections of rAAV2/9-hSyn-DIO-hM4D(Gi)-eGFP-WPRE-pA virus for inactivation of PV interneurons when clozapine N-oxide (CNO) was continuously administered throughout the light flicker treatment period. Saline was used in place of CNO as a control (Gi+Saline group and Gi+Saline+40 Hz group, respectively) (Figure 4J). Both approaches significantly abolished the elevated neurogenesis induced by long-term 40 Hz light flicker treatment, as evidenced by the reduced numbers of BrdU^+^, DCX^+^, and BrdU^+^/DCX^+^ cells in the Gi+CNO+40 Hz group and the Casp3+Saline+40 Hz group (Figures 4K-4N).

Despite extended exposure to 40 Hz light flicker, the quantities of PV interneurons (Figures S3A and S3B) and TBR2^+^ cells (Figures S3C and S3D) remained unchanged, indicating that the light flicker stimulation did not directly impact the proliferation of TBR2^+^ intermediate neuroprogenitor cells in the adult DG. In contrast, deletion and inactivation of PV interneurons significantly reduced the number of TBR2^+^ cells in the DG region (Figure S3E). These results collectively suggest that light flicker-induced adult neurogenesis necessitates the presence of PV interneuron functions, potentially to facilitate the maturation and survival of immature neurons in the DG, rather than to exert a direct effect on the proliferation of neuroprogenitors.

### 40 Hz light flicker enhanced GABA production in the DG region

We next investigated potential alterations in GABA levels within the DG to explore the mechanisms through which PV interneurons support neurogenesis during light flicker treatment.

First, microdialysate samples were collected every 15 minutes, starting 45 minutes before the 1-hour session of 40 Hz light flicker stimulation and continuing for 120 minutes post-treatment (Figure 5A). The GABA levels in the DG were quantified using HPLC. A two-fold increase in GABA levels in the DG occurred following the 1-hour 40 Hz light flicker treatment compared to pre-treatment levels. The GABA levels returned to baseline within 30 minutes after the conclusion of the 40 Hz light flicker treatment.

**Figure 5.**
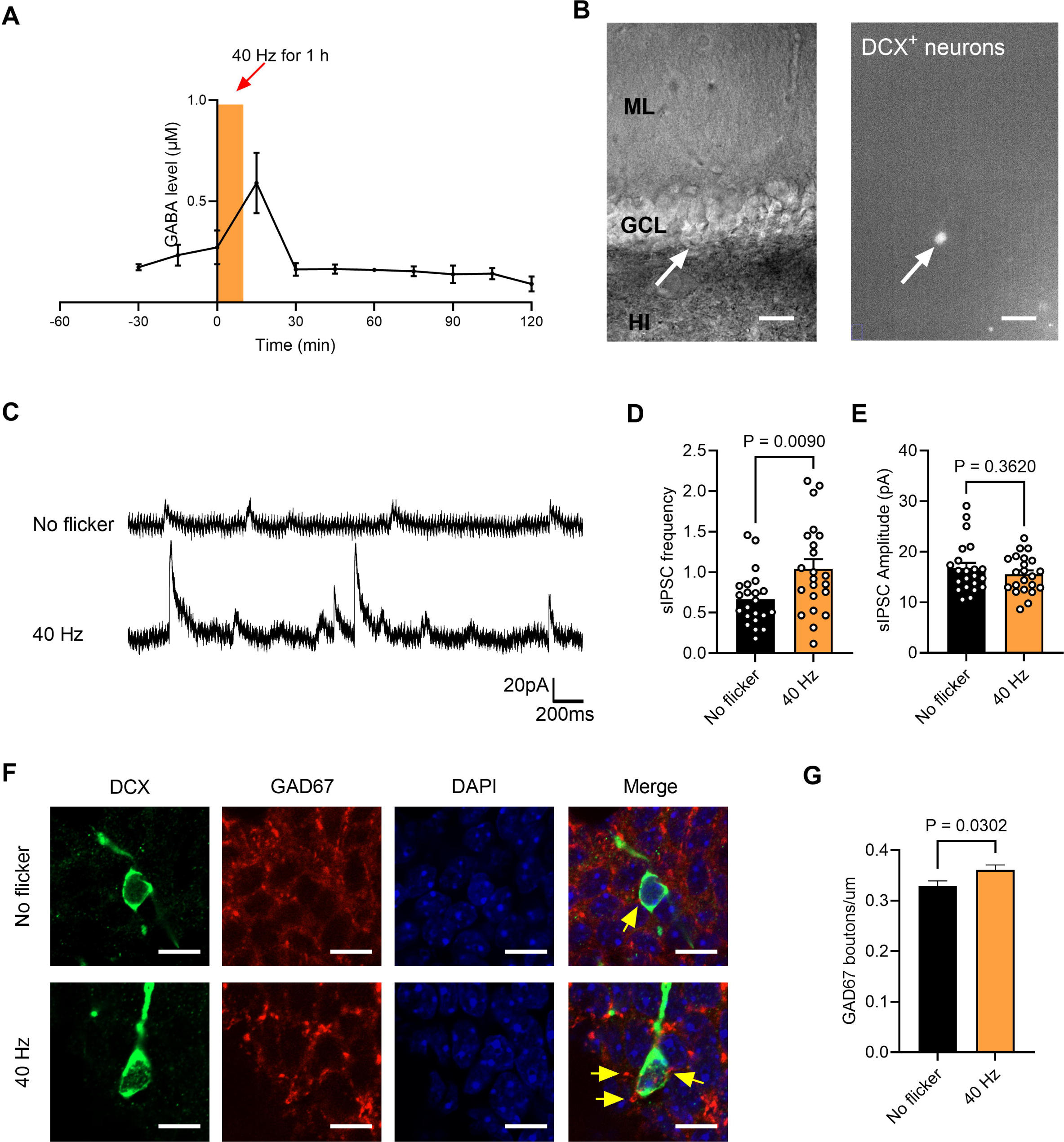
Increased GABA level in the DG region evoked by 40 Hz light flicker. (A) An HPLC analysis of microdialysates from the DG region during and after 1 h 40 Hz light flicker showed an apparent increase in GABA level between 15 – 30 minutes after the treatment. The GABA level returned to an average level afterward. (B) Electrophysiological recordings of DCX^+^ neurons expressing eGFP by infection with rVSVG-Retrovirus-CAG936-eGFP-3xFlag-WRPE-pA. The DCX^+^-eGFP neurons were identified using a microscope under phase contrast bright field (left panel, arrow) and epifluorescent field (right panel, arrow). Scale bars, 20 μm. (C) Examples of recording traces. (D, E) After the 40 Hz light flicker treatment, the IPSC frequency (D), but not the IPSC amplitude (E), of DCX^+^ neurons were increased. (F) Fluorescent confocal microscope images showing the density of GAD67^+^ (red) perisomatic puncta (yellow-colored arrows) on DCX^+^ (green) cells in different groups (see Methods, Scale bars, 20 μm). (G) Significantly increased GAD67^+^ puncta density on the perisomatic region of DCX^+^ cells. Student’s t test was performed for D, E, and G. Specific P values were indicated in the panels.

Second, the electrophysiological characteristics of DCX^+^ neurons were investigated. Mice were injected with rVSVG-Retrovirus-CAG936-eGFP-3xFlag-WRPE-pA in the DG to introduce green fluorescent protein (GFP) expression in newly generated neurons^23^ (Figure 5B). Using patch clamp recording, the sIPSC frequency of these GFP^+^ cells significantly increased, while the sIPSC amplitude remained unchanged, following the 40 Hz light flicker treatment (Figures 5C-5D), indicating that the DCX^+^ cells received enhanced presynaptic inhibition from GABAergic neurons.

Third, the density of GAD67^+^ perisomatic synaptic puncta surrounding DCX^+^ cells increased significantly in mice exposed to 40 Hz light stimulation (Figures 5E-5G). To monitor the temporal variations in GABA levels during the 1-hour 40 Hz light flicker treatment, we conducted an immunohistochemical analysis to assess GABA and GAD67 co-localization in the DG at different time points. These time points comprised no flicker, 15 minutes into the flicker, 30 minutes into the flicker and 30 minutes post-flicker (Figure S4A). The intensity of GABA in the DG exhibited a significant increase within the initial 30 minutes of 40 Hz light flicker, returning to the baseline level observed in the no flicker group 30 minutes post-flicker (Figure S4B). Additionally, the intensity of GAD67 showed a significant increase within the first 30 minutes of 40 Hz light flicker, persisting at a high level even 30 minutes after the conclusion of the 40 Hz light flicker session (Figure S4C).

These data demonstrated enhanced GABA production in response to 40 Hz light flicker treatment.

### Blocking the GABA_A_R in adult mice abolishes enhancement of neurogenesis and spatial learning evoked by 40 Hz stimulation

To investigate the role of GABA in the enhancement of neurogenesis induced by 40 Hz light flicker stimulation, mice were given daily i.p. injections of bicuculline for 2 weeks to block the GABA_A_R (Figure 6A). The sIPSC amplitude of granule cells in the 40 Hz+Bicuculline group showed a significant decrease compared to the 40 Hz+Vehicle group (Figures 6B-6C). In the APA test conducted from day 25 to day 30, mice in the 40 Hz+Vehicle group received significantly fewer shocks on the last 2 testing days than the 40 Hz+Bicuculline group (Figures 6D-6F). Remarkably, the 40 Hz+Bicuculline group mice displayed an inability to learn the location of the shock zone on day 5 of testing. Furthermore, the numbers of BrdU^+^, DCX^+^, and BrdU^+^/DCX^+^ cells in the DG of mice in the 40 Hz+Bicuculline group showed a significant decrease compared to those in the 40 Hz+Vehicle group (Figures 6G-6J). Taken together, blocking GABA_A_R in adult mice eliminated the enhancements in neurogenesis and spatial learning evoked by long-term 40 Hz light treatment.

**Figure 6.**
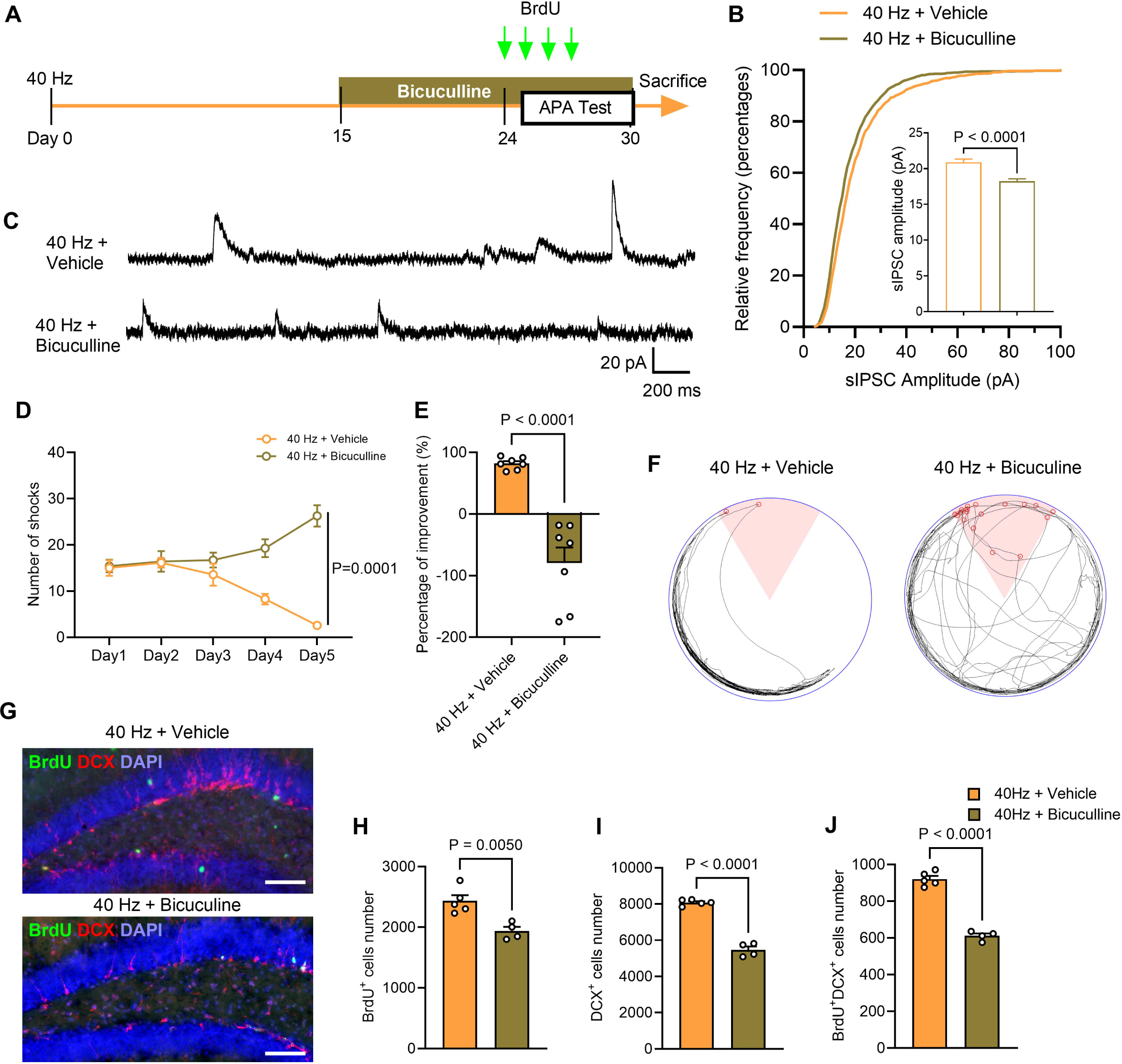
Inhibition of GABA_A_R in adult mice eliminates 40 Hz light flicker-enhanced spatial learning. (A) Experimental schema for GABA_A_R antagonist treatment and APA test. (B-C) Bicuculline treatment significantly decreased the IPSC amplitude of granule cells after 40 Hz light flicker treatment. Examples of recording traces are represented in C. (D) The bicuculline-treated mice received significantly more shocks during the 5 days of APA test, indicating inhibition of enhanced spatial learning by 40 Hz light flicker stimulation. A two-way RM ANOVA with Tukey’s *post hoc* test was performed to demonstrate the significant effect of the 40 Hz light flicker treatment and the bicuculline treatment on shock numbers [F(12, 1) = 30.74, p = 0.0001]. (E) The percentage improvement in avoiding shocks on the 5^th^ day of APA test comparing to the 1^st^ day of APA test. Bicuculline treatment significantly eliminated the improvement of spatial learning conferred by long-term 40 Hz light flicker treatment. (F) The representative movement trajectory map on the 5^th^ day of the APA test. (G) Double immunostaining of BrdU (green) and DCX (red) with DAPI (blue) of the DG region of different groups of mice. Scale bars, 100 μm. (H-J) Bicuculline treatment significantly decreased the number of BrdU^+^, DCX^+^, and BrdU^+^/DCX^+^ cells in the DG after 40 Hz light flicker. Student’s t test was used for panels E, H-J. Specific P values were indicated in the panels. n = 7 mice per group.

## DISCUSSION

The exact mechanism underlying how rhythmic 40 Hz light flicker alleviates neurological functional deficits is currently under intense discussion and investigation^6^. Existing evidence indicates that 40 Hz light flicker entrain with various brain areas, including the hippocampus, to stimulate responses in hippocampal neural circuits^4^, the circadian clock^37^, the glymphatic contraction^3^, microglial activation^1^, and secretion of selective neurochemical mediators such as adenosine^5,7^. However, these studies have primarily focused on acute and short-term exposure to 40 Hz light flicker. The current study represents the first comprehensive analysis of long-term treatment with 40 Hz light flicker and presents compelling evidence that prolonged exposure to 40 Hz light flicker promotes robust neurogenesis in the DG region of the adult brain and enhances spatial learning. Our data showed that these mice did not display increased levels of stress and anxiety, therefore supporting the continued exploration of this relatively simple technique as a potential clinical therapy for treating neurological diseases in humans.

The long-term beneficial effect of 40 Hz light flicker was dependent on DG neurogenesis. This observation was substantiated by the reversal of spatial learning enhancement following the elimination of newly formed DCX^+^ immature neurons in the DG region using adult DCX^DTR^ transgenic mice (Figure 2). Adult neurogenesis occurs within a complex local environment called the neurogenic niche, which supports neural precursor cells during their maturation. The 40 Hz light flicker specifically increased neurogenesis in the DG but not in the SVZ (Figure 1). The underlying reason for this discrepancy remains unclear. We suspect that the neurogenic niche in the SVZ may not respond to the 40 Hz light flicker stimulation.

A key feature of adult neurogenesis is its dynamic regulation by neuronal activity during specific stages^24,38^. For instance, voluntary exercise (wheel running, RUN) and environmental enrichment (ENR) led to a doubling in the production of new hippocampal granule cells while not impacting the number of adult-generated olfactory granule cells^10,11,13,35,39^. Although both RUN and ENR boost adult hippocampal neurogenesis, they operate through different mechanisms^40^. RUN stimulates the proliferation of precursor cells, whereas ENR enhances adult hippocampal neurogenesis by boosting the survival and maturation of newly generated neurons^13,39^. In this context, the 40 Hz light flicker’s mechanism of enhancing neurogenesis aligns more closely with ENR. Several pieces of experimental data support this assertion. First, after the long-term stimulation, there were no changes in DG microcirculation, such as the diameter of capillaries and the regional blood flow (Figure S2). Our previous studies also demonstrated no regional cerebral blood flow changes during short-term 40 Hz light flicker treatment^4^. Alteration of cerebral blood flow is a significant factor contributing to physical exercise RUN-induced cell proliferation in the DG, while ENR does not affect cerebral blood flow^40,41^. Second, the ablation of PV interneurons significantly decreased the number of TBR2^+^ cells in the DG, affirming the necessity of PV-dependent proliferation. However, long-term exposure to 40 Hz light flicker did not increase the number of TBR2^+^ cells in the DG region (Figure S3). In contrast, reducing GABA levels by deleting and inactivating GABAergic PV interneurons or directly applying a GABA_A_R antagonist significantly decreased the number of DCX^+^ cells. Together, this data indicates that the 40 Hz light flicker plays a crucial role in supporting the maturation of DCX^+^ immature granule cells through the GABAergic inhibitory functions of PV interneurons in a mechanism similar to ENR.

Multiple stages of adult neurogenesis are precisely regulated by PV interneurons and their GABAergic transmission^42^. Contrary to expectations, long-term treatment with 40 Hz light flicker did not increase the number of PV interneurons in the DG. However, our study provides compelling evidence demonstrating that 40 Hz light flicker activated PV interneurons and upregulated GABA levels. For example, fibre photometry data revealed significantly increased calcium signaling in specifically labeled PV interneurons activated by 40 Hz light flicker (Figure 3). Following 30 days of light flicker treatment, the excitability of PV interneurons increased, as evidenced by *in vitro* electrophysiology results (Figure 3), suggesting alterations in membrane channels of PV interneurons.

Furthermore, eliminating PV interneurons through caspase-3 expression (Figure 4) significantly dampened gamma oscillation in the DG, as gamma rhythmic flicker primarily synchronizes with fast-spiking PV interneurons. Inhibiting PV interneurons using an inhibitory rAAV-Gi virus markedly abolished the elevated level of neurogenesis evoked by prolonged exposure to 40 Hz light flicker. These findings unequivocally demonstrated that PV interneurons were activated during extended 40 Hz light flicker treatment and its activity was required for neurogenesis and the enhancement of cognitive function.

GABA serves as a primary extrinsic factor that governs the differentiation of Type-2 intermediate neuroprogenitor cells within the DG region through GABA_A_R^43^. The GABA_A_R agonist significantly increases the number of new neurons labeled with BrdU^33,42^. Indeed, following the 40 Hz light flicker treatment, there was a notable elevation in GABA levels within the DG, substantiated by microdialysis and HPLC analysis. Inhibitory GABAergic synaptic connections to the newly generated neurons were also enhanced, evidenced by an increased presence of GAD67^+^ buttons surrounding granule cells co-localizing with DCX in the DG region (Figure 5). Significantly, blocking GABA_A_R inhibitory transmission using bicuculline mitigated the improvements in spatial learning and neurogenesis in the DG region induced by the 40 Hz light flicker (Figure 6).

Together, these findings demonstrated that neurogenesis induced by 40 Hz light flicker involves a mechanism that relies on the activity of PV interneurons and GABA tonic support for the postmitotic immature neurons in the adult DG to promote their maturation and synaptic integration. The current study provided new evidence demonstrating the long-term beneficial effects of rhythmic light flicker as a potentially effective treatment for neurological diseases.

## SUPPLEMENTAL INFORMATION

**Supplemental information can be found online at XXX**

## STAR ★ METHODS

Detailed methods are provided in the online version of this paper and include the following:

- KEY RESOURCES TABLE
- RESOURCE AVAILABILITY

- Lead contact
- Materials availability
- Data and code availability
- EXPERIMENTAL MODEL AND ANIMAL STDY ETHICAL APPROVAL

- Experimental animals
- Ethical approval and Animal experimentation design
- METHOD DETAILS

- Drug administration
- Visual stimulation protocol
- Open field test
- Active place avoidance (APA) test
- Chemogenetic inhibition of PV interneurons
- Stereotaxic surgery
- Brain slice electrophysiology
- Quantification of neurogenesis by immunofluorescent staining
- Analysis of GAD67 perisomatic puncta
- Fiber photometry recording and analysis
- Microdialysis and HPLC analysis
- Recordings of local field potential (LFP)
- Insertion of GRIN lens in hippocampus DG
- Microvasculature imaging using two-photon microscopy
- QUANTIFICATION AND STATISTICAL ANALYSIS

## Acknowledgments

We thank the SUSTech Animal Facility for animal maintenance and SUSTech Core Research Facilities for imaging acquisition supports. Financial support for JJ and STH was from grants from the National Natural Science Foundation of China (82301730, 32371029, respectively). STH was supported by grants from the Shenzhen-Hong Kong Institute of Brain Science-Shenzhen Fundamental Research Institutions (2023SHIBS0002), and Shenzhen Medical Research Fund (B2301001). STH is also a member of the Guangdong Innovation Platform of Translational Research for Cerebrovascular Diseases. STH, TW and PB are members of the SUSTech-UQ Joint Centre for Neuroscience and Neural Engineering (CNNE). STH is a Pengcheng Peacock Plan (A) Distinguished Professor.

## Author Contributions

Hai Yan, Xufan Deng, Mei Yu, Shiyu Wu and Yunxuan Wang did the behaviour tests. Hai Yan, Xufan Deng, Shiyu Wu, Yunxuan Wang, Jingyuan Mai, Yifan Pan, Jinghong Chen and Yunxuan Wang did the immunostaining. Jun Ju, Yifan Pan and Jinghong Chen performed electrophysiological experiments. Shiyu Wu, Bo Liu, Huimei Wang and Zhengyu Zhang did the local field potential recording and analysis. Jun Du and Shiyu Wu did the microdialysis sample collection, and Jun Du did HPLC analysis. Shiyu Wu did the fiber photometry recording and analysis. Perry Bartlett and Tara Walker provided the DCX inducible deletion mice. Sheng-Tao Hou, Jun Ju, and Yizheng Wang designed the project, analysed data, supervised the project and wrote the draft. All authors contributed to the article and approved the submitted version.

## STAR ★ METHODS

### KEY RESOURCES TABLE

| REAGENT OR RESOURCE | SOURCE | IDENTIFIER |
| --- | --- | --- |
| **Experimental organisms** |  |  |
| Mouse: C57BL/6J | Vital River Lab Animal Technology (Zhejiang, China) | Cat# 213 |
| Mouse: DCX<sup>DTR</sup> | Queensland Brain Institute | Refence 15 |
| Mouse: PV-Cre (B6.129P2-*Pvalb*<sup>tm1(cre)Arbr</sup>/J) | The Jackson Lab, USA | Cat# 017320 |
| **Antibodies** |  |  |
| Anti-BrdU antibody | Abcam | Ab6326 |
| Anti-DCX antibody | Abcam | Ab18723 |
| Anti-GABA antibody | Sigma-Aldrich | A2052 |
| Anti-GAD67 antibody | Abcam | Ab26116 |
| Anti-TBR2 antibody | Abcam | Ab23345 |
| Anti-PV antibody | Abcam | Ab181086 |
| **Chemicals** |  |  |
| 5'-Bromo-2'-deoxyuridine (BrdU) | Sigma-Aldrich | B5002 |
| Diphtheria Toxin (DT) | Sigma-Aldrich | D0564 |
| Clozapine N-oxide (CNO) | MedChemExpress | HY-17366 |
| Bicuculline | MedChemExpress | HY-N0219 |
| DAPI | Abcam | Ab104139 |
| Goat serum | Gibco | 16210072 |
| Alexa Fluor<sup>TM</sup> 647-Hydrazide | Thermo Fisher Scientific | A20502 |
| FITC-dextran (70kDa) | Sigma-Aldrich | 46945 |
| CNQX | Sigma-Aldrich | C239 |
| AP5 | Sigma-Aldrich | A8054 |
| **Recombinant viral strains** |  |  |
| rAAV2/9-hSyn-DIO-hM4D(Gi)-eGFP-WPRE-pA | Shanghai Taitool Bioscience Co.Ltd., China | S0286-9 |
| rVSVG-Retrovirus-CAG936-eGFP-3xFlag-WRPE-pA | Shanghai Taitool Bioscience Co.Ltd., China | VSVG-R5010 |
| rAAV2/9-hSyn-DIO-jGCaMP7f-WPRE-pA | Shanghai Taitool Bioscience Co.Ltd., China | S0588-9 |
| rAAV2/9-CAG-DIO-taCaspase3-TEVp-WPRE-pA | Shanghai Taitool Bioscience Co.Ltd., China | S0236-9 |
| **Software and algorithms** |  |  |
| GraphPad Prism | Graphpad | Version 10 |
| Ethovision XT | Noldus | Version 15 |
| FIJI | ImageJ | Version 1.54i |

### RESOURCE AVAILABILITY

#### Lead contact

Further information and requests for resources and reagents should be directed to and will be fulfilled by the Lead Contact, Sheng-Tao Hou.

#### Materials availability

This study did not generate new unique reagents.

#### Data and code availability

- All data reported in this paper will be shared by the lead contact upon request.
- This paper does not report the original code.
- Any additional information required to reanalyze the data reported in this paper is available from the lead contact upon request.

### EXPERIMENTAL MODEL AND ANIMAL STDY ETHICAL APPROVAL

#### Experimental animals

C57BL/6J mice were purchased from the Vital River Laboratory Animal Technology Co., Ltd. (Zhejiang, China). PV^Cre^ Mice (B6.129P2-*Pvalb*^tm^^1(cre)Arbr^/J) were purchased from Jackson Laboratory (Strain # 017320, USA) and bred locally. To ablate DCX-positive (DCX^+^) neurons, a knock-in mouse model in which the diphtheria toxin (DT) receptor (DTR) was expressed under the control of the endogenous DCX promoter sequence via an internal ribosome entry site sequence was generated and used as previously described^44^. In these mice, DTR is expressed under the control of the DCX promoter, which allows for specific ablation of immature DCX-expressing neurons after administration of DT while leaving the neural precursor pool intact. DCX^DTR^ knock-in mice were bred on a C57BL/6J background and backcrossed for more than seven generations.

Male mice at the age of 6 months old were used for the study. Animals were maintained in a condition-controlled room in a pathogen-free SPFII animal facility (23 ± 1 °C, 50 ± 10% humidity). A 12:12 h light/dark cycle (7 a.m. to 7 p.m.) was automatically imposed, and the light intensity was maintained at 15-20 lx during the light period. However, the animal facility room light was at 200 lx during cleaning and experimental operation. Mice were housed in groups of six per individually ventilated cages and given access to food and water *ad libitum*. Experimenters were blinded to animals’ treatments and sample processing throughout the subsequent experimentation and analyses.

#### Ethical approval and Animal experimentation design

Animal experiment protocols were approved by the Animal Care Committee of the Southern University of Science and Technology (Shenzhen, China). The ARRIVE guideline was followed when designing, performing, and reporting animal experimentation^45^. Mice used in the current study were randomly assigned to each group to maintain total randomization.

According to the ARRIVE reporting guidelines, efforts were made to minimize the number of animals and animal suffering. The inclusion criterion was based on the pre-established identical age and sex of the mice. AEEC Animal Experimentation Sample Size Calculator was used to determine the minimum sample size required to test the study hypothesis^46^. Results indicated the required sample size to achieve 80% power for detecting 25% difference between two independent means at a significance criterion of α = 0.05 was n = 3. A minimum of six mice per group were used in the behavioral studies to achieve meaningful statistical differences. At least three mice per group were used for brain slice electrophysiology, immunostaining, fiber photometry recording, microdialysis, and local field potential recording.

### METHOD DETAILS

#### Drug administration

Saline was used to make 10 mg/ml 5’-Bromo-2’-deoxyuridine (BrdU; B5002; Sigma) and 0.3 mg/ml clozapine N-oxide (CNO; HY-17366; MedChemExpress). The intraperitoneal (i.p.) administrations of BrdU (100 mg/kg) were carried out for the first four days of the last week of long-term 40 Hz light flicker treatment and CNO (3 mg/kg) was carried out for 30 days during 40 Hz light flicker treatment. Diphtheria Toxin (DT; D0564; Sigma) was dissolved in phosphate-buffered saline (PBS). The i.p. injection of DT (10 μg/kg) was carried out every 2 days for 2 weeks from the second week of 40 Hz light flicker treatment. Bicuculline (0.05 mg/ml, HY-N0219, MCE) was dissolved in saline containing 1% DMSO and was injected i.p. at 0.5 mg/kg for 2 weeks during the 40 Hz light flicker treatment.

#### Visual stimulation protocol

The visual stimulation paradigm and methods were essentially as previously described^4^. The visual stimulation equipment, TangGuang^TM^, comprised seven modules, which included a tunable frequency signal generator and six LED lamps (36 V). The six LED lamps were interconnected in parallel circuits and evenly positioned around two transparent cages housing the control and experimental groups simultaneously. The light stimulation was administered consistently once daily from 6 to 7 pm over the course of the experiment, lasting up to 30 days.

#### Active place avoidance (APA) test

APA test is an effective, versatile, and repeatable tool to test hippocampus-dependent spatial learning in rodents. The testing protocol was precisely as previously described^47^. The mice were placed on a gridded platform with a 1-meter diameter, enclosed by a clear Perspex cylinder (height: 32 cm; diameter: 77 cm) that rotated at 1 revolution per minute (Bio-Signal Group, USA). A tracking computer defined a 60° shock zone, where any animal entering this region would receive a 0.5 mA foot shock after a 0.5-second delay, with an inter-shock delay of 1.5 seconds. Four visual cues were evenly distributed around the room, and the animal was consistently positioned in the arena opposite the shock zone for each day of testing.

The APA test protocol included one day of habituation without shocks for 5 minutes, followed by 5 days of testing with shocks administered for 10 minutes each day. The grid and platform were cleaned using 75% ethanol between testing trials. Parameters such as total distance traveled, number of shocks received, and number of entries into the shock zone were automatically recorded using a ceiling-mounted video camera positioned overhead. The learning ability during the testing period was evaluated by comparing the parameters recorded on the 1^st^ day with those on the 5^th^ day of testing for each animal. The performance improvement was calculated as a percentage, reflecting the progress made during the task.

#### Chemogenetic inhibition of PV interneurons

To selectively inhibit PV interneurons, PV^Cre^ mice received a stereotactic injection of rAAV-DIO-hM4D(Gi)-eGFP (serotype 9, 120 nl) into the DG hilus. The vector control AAV2/9-hSyn-DIO-EGFP was utilized for comparison. All AAV viruses were procured from Taitool Bioscience Co. Ltd (Shanghai, China) and were diluted with PBS to a final concentration ranging between 5×10^12 and 1×10^13 genome copies per ml before stereotaxic administration into the mouse brain. Following a two-week interval, mice were treated with 3 mg/kg of CNO via i.p. injection daily throughout the entire light flicker treatment period (30 days). Control mice received equivalent volumes of saline solution. Subsequently, the mice underwent APA test and neurogenesis analysis.

#### Stereotaxic surgery

Mice were anesthetized with isoflurane and then received bilateral injections of 200 nl of VSVG-Retrovirus-CAG936-eGFP-3xFlag-WRPE-pA (concentration at 1.34E + 9 v.g./ml; v.g. viral genome). PV^Cre^ mice were also anesthetized with isoflurane and bilaterally injected with 200 nl of AAV2/9-CAG-DIO-taCaspase3-TEVp-WPRE-pA (concentration at 3.3E + 12 v.g./ml) or AAV2/9-hSyn-DIO-hM4D(Gi)-eGFP-WPRE-pA (concentration at 3.3E + 12 v.g/ml) into the DG at a rate of 0.05 µl/min. The injection site was precisely located at AP: -2.0 mm, ML: ±1.5 mm, DV: -2.2 mm from bregma. Following injection, the needle was held in place for 5 minutes before being removed, and the mice were allowed to recover in their home cage until fully awake. To manage discomfort, meloxicam (1 mg/kg, s.c.) and penicillin (3000 U per mouse, i.p.) were administered once daily for three days.

#### Brain slice electrophysiology

The protocol was modified from our previous studies^48,49^. Mice were anesthetized with 1% pentobarbital sodium and decapitated. Mouse brain was dissected out and immersed in ice-cold artificial CSF (aCSF) containing (in mM): 30 NaCl, 26 NaHCO_3_, 10 D-glucose, 4.5 KCl, 1.2 NaH_2_PO_4_, 1 MgCl_2_, 194 sucrose, adding 1.5 mL 1M HCl per 1 L cutting solution and bubbled with 95% O_2_/5% CO_2_. Coronal brain slices (350 μm) were made on a vibratome (VT1120S, Leica Systems). Slices were allowed to recover for 30 min at 34℃ in aCSF containing (in mM): 124 NaCl, 26 NaHCO_3_, 10 D-glucose, 4.5 KCl, 1.2 NaH_2_PO_4_, 1 MgCl_2_, 2 CaCl_2_, adding 10 g sucrose and 1 mL 1M HCl per 1 L aCSF and bubbled with 95% O_2_/5% CO_2_. After being transferred to a holding chamber at room temperature, the recording started only after at least one hour of recovery. The slices were then placed in a recording chamber (RC26G, Warner Instruments, USA) on the x-y stage of an upright microscope (BX51W; Olympus, Tokyo, Japan) and were perfused with aCSF at a rate of 2 ml/min. All recordings were conducted at room temperature.

To record spontaneous IPSC (sIPSC) in DCX^+^ neurons, patch recording pipettes were filled with (in mM) 125 CsMeSO_3_, 5 NaCl, 10 HEPES (Na^+^ salt), 5 QX314, 1.1 EGTA, 4 ATP (Mg^2+^ salt), 0.3 GTP (Na^+^ salt). sIPSC was recorded in aCSF adding 50 μM D(-)-2-Amino-5-phosphonopentanoic acid (AP5) and 10 μM CNQX at +10 mV. The acquisition frequency was 20.0 kHz, and the filter was set to 2.9 kHz.

To examine PV interneuron excitability, recording pipettes were filled with (in mM): 128 potassium gluconate, 10 NaCl, 10 HEPES, 0.5 EGTA, 2 MgCl_2_, 4 Na_2_ATP, and 0.4 NaGTP. Current steps ranging from 0 pA to 300 pA with a 20 pA increment and 1-second duration were utilized. The increment was later adjusted to 5 pA to determine the rheobase current, which was defined as the minimum current required to elicit an action potential. The resting membrane potential was measured by injecting a current of 0 pA. The acquisition frequency was set at 20.0 kHz, and the filter was set to 2.9 kHz. sIPSC were analyzed using Mini-analysis (Synaptosoft Inc.). The frequency and amplitude were analyzed. In addition, neuronal spiking traces were imported into Fitmaster software (HEKA Elektronik), and then the number of action potentials was measured.

#### Quantification of neurogenesis by immunofluorescent staining

The immunostaining protocol was exactly as we previously described^4,50–52^. Briefly, mice were administered phenobarbital sodium salt (0.1 g/kg) to induce anesthesia. Subsequently, the mice were subjected to transcardial perfusion with ice-cold 0.01 M PBS, followed by 4% (weight/volume) paraformaldehyde (PFA) in 0.01 M PBS. Then the brains were retrieved and dehydrated in 15% sucrose for 1 day, and 30% sucrose for 2 days. Serial coronal brain sections (40 μm thickness) through the hippocampus were cut and retained for immunostaining. A total of 72 consecutive slices from the beginning of the DG (at AP axis = -1.4mm from bregma, coronal view) were collected and then preserved in cryo-protectant solution at -20°C. One-sixth of each brain was pooled into one series for immunohistochemistry.

Brain slices were retrieved from cryo-protectant solution and washed a float 3 times in PBST before staining. The slices were transferred onto glass slides and incubated in 0.5% Triton X-100 at 37°C for 30 min. After permeabilization and washing with PBST, the slices were heated for 2 minutes in citrate buffer (pH = 6.0) with an autoclave (105 °C) for antigen retrieval. After retrieval, the slices were cooled down in citrate buffer to room temperature, followed by PBST washing. For BrdU immunostaining, sections were treated with pre-warmed 1 M HCl for 30 min at 37°C to disrupt hydrogen bonds between bases and to denature the DNA. These slices were then blocked in 10% goat serum PBS with 0.2% TritonX-100 at room temperature for 1h. The primary antibody binding was carried out in 3% goat serum PBST solution with 0.2% Triton X-100 at 4°C overnight. The following primary antibodies were used: anti-BrdU (Abcam, rat, 1:500), anti-DCX (Abcam, rabbit, 1:500), anti-GABA (Sigma, rabbit, 1:500), anti-GAD67 (Abcam, rabbit, 1:500), anti-TBR2 (Abcam, rabbit, 1:500), and anti-PV (Abcam, rabbit, 1:500). On the second day, the slices were washed with PBST 3 times and incubated with the secondary antibody at room temperature for 1h. After washing, the slides were observed using a Zeiss LSM980 confocal microscope.

The methodology employed to quantitatively assess the quantities of DCX and BrdU-positive cells for neurogenesis was as outlined in a previous study^41,52,53^. Positive cells were counted on every sixth section (240 μm apart) across the entire dorsal-ventral span of the DG. The counts were then multiplied by six to determine the total number of labeled cells within the DG of each brain.

#### Analysis of GAD67 perisomatic puncta

Images were captured using a Zeiss LSM980 confocal microscope equipped with a 63× lens. We obtained 3-4 sections from each mouse for analysis. To analyze perisomatic puncta in the images, the cell soma profile was manually outlined in ImageJ software, and a series of custom-made macros in Fiji were used, as previously described. To define the region of interest (ROI), the initial outline was expanded by 1 µm from the cell body’s surface. The ROI was established as the enclosed area between these two outlines. Subsequently, the total number of puncta within the ROI was counted, and the perimeter of the cell soma was calculated. The density of GAD67 perisomatic puncta was determined by dividing the number of puncta by the perimeter^54,55^.

#### Fiber photometry recording and analysis

The mice were anesthetized with isoflurane and received a unilateral injection of 250 nl of AAV2/9-hSyn-DIO-jGCaMP7f-WPRE-pA (Taitool, China) into the DG at a rate of 0.05 µl/min. The injection site was targeted at AP: -2.0 mm, ML: ±1.5 mm, DV: -2.2 mm from bregma. The needle was left in place for 10 minutes post-injection before being removed. Subsequently, the mice were allowed to recover in their home cages until fully awake. To alleviate post-operative pain, meloxicam (1 mg/kg, s.c.) and penicillin (3000 U per mouse, i.p.) were administered once daily for 3 days.

After a three-week period, to capture the fluorescence emitted by the calcium sensor, an optical fiber was attached to the implanted ferrule via a ceramic sleeve. A commercial fiber photometry system (Thinker Tech Nanjing Biotech, China) was employed to record the emissions. The recordings were carried out on freely moving mice during light flicker visual stimulation. The fiber photometry data underwent initial processing using the pMat application (https://github.com/djamesbarker/pMAT) to handle raw data from individual trials^56^. In the heatmaps, the signals were normalized within the range of -5 to 10 for each trial. Prior to further analysis, the raw signals were corrected to address any photo-bleaching effects by accounting for the overall trend. Fluorescence signals obtained from the GCaMP channel were assessed and normalized to quantify fluctuations in fluorescence (normalized ΔF/F). Subsequently, the signals were transformed into a robust z-score utilizing the following formula:

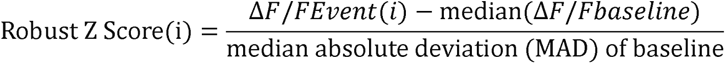

To conduct additional analysis, peri-event time histograms (PETH) were created by aggregating data from multiple trials within time windows corresponding to 40 Hz light flicker treatments. The area under the curve was calculated for the segment within each user-defined window using the trapezoidal method for integral computation (employing the MATLAB *trapz* function).

To determine the intensity of the calcium signal to 40 Hz light flicker visual stimulation, ΔF/F0 values were calculated, a stable baseline was established, and the standard deviation (SD) of the baseline was determined. A ΔF/F0 value exceeding 3 times the SD, and meeting the automatic multiscale-based peak detection algorithm criteria^4,52^, was considered a genuine calcium signal rather than background noise. The area under the curve was computed for the segment falling within each user-defined window using the trapezoidal method for integral calculation (utilizing the MATLAB *trapz* function).

#### Microdialysis and HPLC analysis

The procedures used were precisely as previously described^57^. An MBR intracerebral guide cannula (MD-2255, BASI) was surgically implanted into the DG of the mice at coordinates AP: -2.0 mm, ML: ±1.5 mm, DV: -2.2 mm from bregma. Prior to the experiment, the mice were anesthetized, and an MBR-1-5 brain microdialysis probe (MD-2211, BASI) was inserted into the cannula. Artificial cerebrospinal fluid (aCSF) containing the following components in mM: 124 NaCl, 26 NaHCO_3_, 10 D-glucose, 4.5 KCl, 1.2 NaH_2_PO_4_, 1 MgCl_2_, 2 CaCl_2_, supplemented with 10 g sucrose and 1 mL 1M HCl per 1 L aCSF, and bubbled with 95% O_2_/5% CO_2_, was infused into the microdialysis system at a constant rate of 1 μl/min using a syringe pump. The dialysates were continuously collected for 60 minutes under stable lighting conditions. Subsequently, following 1 hour of exposure to 40 Hz light flicker, the collection period was extended to 60 minutes, with dialysates collected every 15 minutes while the mice remained under anesthesia. The concentration of adenosine in the dialysates was then quantified using HPLC analysis.

#### Recordings of local field potential (LFP)

The procedures used were as precisely described^4^. *Implantation of electrodes*: Mice were anesthetized with a mixture of 2.5% isoflurane in N_2_:O_2_ (70:30; flow rate 400 ml/min) in an induction chamber. Anesthesia was maintained with 1.5% isoflurane in N_2_:O_2_ mixture during surgery using an isoflurane vaporizer (#R540, RWD Life Science, Shenzhen, China). The animal’s body temperature was maintained at 37.0 ± 0.5 °C using a rectal temperature probe and a heating pad (#TCAT-2DF, Harvard Apparatus, USA).

A cranial window at 1.2 mm in diameter with AP: -2.0 mm, ML: ±1.5 mm, DV: -2.2 mm from bregma over the DG was created using a dental drill (#78001, RWD Life Science) guided by stereotaxis (# 68861 N, RWD Life Science). A 4-channel microwire array electrode (35 μm, Stablohm 650, California Fine Wire Co., USA) was inserted into DG and immobilized using kwiksil (Item #KWIK-SIL, Microprobes for Life Science, Gaithersburg, MD, USA). The tetrode was anchored to the skull bone using four skull screws (1.0 mm diameter) and embedded in dental cement. Mice were moved into a warm (37.0 ± 1 °C) recovery chamber (#DW-1, Harvard Apparatus) for 1 h.

*LFP recording and analysis:* After a 4-day recovery period following electrode implantation surgery, the mice were acclimated to the recording setup by being placed in a box measuring 50 cm in depth, 50 cm in width, and 50 cm in length for 10 minutes per day until the first recording session. Prior to each recording session, the box was sanitized using 75% ethanol. During the recording session, a helium balloon delicately supported the implant, enabling the mouse to move freely within the box.

The head stage was connected to the OmniPlex Neural Recording Data Acquisition System (Plexon Inc, Dallas, TX). A camera was set in the front of the box to record the behavioral states of the mouse. An LED lamp was put in front of the box to generate different frequencies of stimulation lights. LFPs were recorded for at least 20 min for each mouse. The LFP signals were sampled at 1000 Hz with a band-pass filter set at 0.5 - 120 Hz. Raw data were stored for later offline analysis. A total of five mice in each group were used for statistical analysis.

Concurrent with data recording, we also recorded the behavioral activities of the mice. The specific time intervals were determined by observing the video of the mice’s behavior, and several epochs were selected from the original data, each with the same length of time.

A multitaper fast Fourier transform method was implemented for power spectral analyses using MATLAB (ver 9.11.0, R2021b) software. Data was filtered with a band-pass filter of 0.1 to 100 Hz and a notch filter of 50 Hz. The power spectrum in Figure 4 was given by a multitaper estimation method using MATLAB using Equation (1). Given a time series *X_n_*, *n* = 1, 2, …, *N*, the number of the Slepian sequences is *K = 2NW−1*. The simplest multitaper estimate of the spectrum is given by

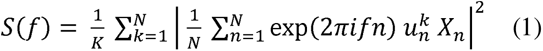

where 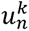, n = 1, 2, …, N is the *k*th Slepian sequence.

The Short-time Fourier transform (STFT) was applied to reveal the power changes in different frequencies on a time scale. The LFP data was segmented into epochs of the same time interval as described above, and a hamming window with a length of 8000, 50% of overlap was used to calculate the spectral amplitude between 1 and 100 Hz through a MATLAB (Ver 9.11.0, R2021b) function *stft*.

#### Insertion of GRIN lens in hippocampus DG

The mice were anesthetized using a gas anesthesia machine (RWD Life Science) and then carefully positioned on a stereotaxic apparatus. A stereoscope was adjusted to provide a clear view of the surface of the mouse skull. Eye ointment (Guangzhou Pharmaceutical Holdings, China) was gently applied, and a heating pad set to 35°C (Physitemp, USA) was utilized for maintaining body temperature. Following this, the hair and scalp were meticulously removed with sterilized scissors, and the subcutaneous tissue was cleaned using a cotton swab.

To ensure precision, the heights of bregma and lambda were aligned, along with lateral heights. Using a cranial drill (RWD Life Science), two 1.0 mm diameter holes were created in the frontal bone and secured with cranial pins from the same manufacturer. Subsequently, with the bregma as the reference point, coordinates were set at ML: -1.5 mm, AP: -2 mm, and a 1.2 mm diameter hole was drilled at this location. The skull fragment was removed, and careful aspiration was performed to expose the hippocampal fibrous capsule and the molecular layer. Additional aspiration was done carefully, and any bleeding was stopped with a hemostatic sponge (Guilin Fukangsen Medical Device, China).

The GRIN lens was gently inserted into the hole while activating the imaging mode of the nVoke imaging system (Inscopix Inc., USA) until the lens bottom was visible. The lens was secured using bio-compatible silicon gel (WPI Surgical Instruments, USA), and a titanium ring was affixed around the lens with acrylate adhesive. Dental cement (Shanghai Yuyan Instruments, China) was applied to cover exposed areas of the skull, with a drop of silicon gel on the lens surface, for additional protection against damage.

#### Microvasculature imaging using two-photon microscopy

Mice equipped with the GRIN lens were anesthetized using a gas anesthesia machine (RWD Life Science). Their hindlimb was secured with sticky tape, and hair was removed using depilatory cream. The skin was sanitized with iodine tincture. A 200 µl mixture comprising 150 µl of 70 kDa Dextran-FITC and 50 µl of Hydrazide-Alexafluor 647 (Thermo Fisher, USA) was slowly injected using an insulin injector after cutting open the skin. Bleeding was controlled with a hemostatic sponge, and the skin was sealed using a suture needle and thread (Shanghai Jinhuan Medical, China). The silicon gel previously applied to the top of the GRIN lens was removed using forceps. The cranial window or GRIN lens surface was cleaned with an alcohol-soaked hemostatic sponge. The mice were then fixed onto a custom-made module through the implanted titanium ring to ensure their position under the objective. Imaging commenced once the mice had regained consciousness.

An upright FVMPE-RS multiphoton microscopy system (Olympus, Japan), based on an Olympus upright microscope equipped with a water objective (25X/1.05, W.D. = 2 mm), was activated. A femtosecond-pulsed Ti:Sa laser (Mai Tai DeepSee, Spectra-physics, USA) was utilized to generate a 1000 nm laser for the excitation of both FITC and Alexa Fluor 647. The laser wavelength was set at 1000 nm, with a power of 15% of 1.28W. Two acquiring channels, FV30-FGR (495-540 nm) and FV30-FRCY5 (660-750 nm), were configured. The image format was adjusted to 512x512, digital magnification to 3X (resulting in a pixel size of 0.931 µm/pixel), and the scanning mode was set to Galvo and unidirectional.

The module’s position under the objective was fine-tuned until the image was visible in the visual field. Imaging of the mice was conducted without 40 Hz light flicker in both the visual cortex and DG. The imaging of the DG was repeated the next day without light flicker. An aluminum foil was affixed to the mice’s head to separate the lens and eyes. A customized 40 Hz light flicker LED through a conducting tube focused light exclusively on the eyeball during imaging.

The Hydrazide signal obtained through the CY5 channel indicated the arteriole lumen. The location where the hydrazide signal ends along the vasculature was designated as the transition point between arteriole and capillary. The 1^st^ to 4^th^ branched capillaries counted from the transition point were classified as pre-capillary, while the 5^th^ to 8^th^ branched capillaries were termed post-capillary. The last segment of the arteriole with a Hydrazide signal situated just before the transition point was labeled the terminal arteriole. The FITC signal captured via the FGR channel depicted the flow of blood cells through the vessel. Each vessel was imaged to showcase its morphology and length. Three-line scans were conducted by aligning the scanning line in the vessel’s extending direction, with a scanning time restricted to 1 second. Between 3 to 8 vessels from each defined category in arteriole, pre-capillary, and post-capillary were randomly imaged across the entire visual field. After imaging, the mice were removed from the module and returned to their rearing cage. Data was collected for subsequent analysis.

### QUANTIFICATION AND STATISTICAL ANALYSIS

All data were represented as the Mean ± standard error of the Mean (SEM). All analyses were performed using Prism (V10, GraphPad Software, La Jolla, CA, USA). Data distribution was determined using the Shapiro-Wilk test first before applying an appropriate parametric or nonparametric statistical test. If the data from the two groups passed the normality test, an unpaired t-test was used. Otherwise, the Mann-Whitney U test was applied. If the data from several groups passed the normality test, One-way ANOVA or two-way ANOVA with Tukey’s *post hoc* test was used. Otherwise, the Kruskal-Wallis test with Dunn’s *post hoc* test was applied. The firing rate was analyzed using two-way RM ANOVA with Tukey’s *post hoc* test or Sidak’s *post hoc* test. Statistical details for specific experiments, including the exact n values and what n represents, precision measures, statistical tests used, and the definitions of significance, can be found in figure legends. A P < 0.05 was considered statistically significant.

