## Supplemental Figures for "Enhancement of spatial learning by 40 Hz visual stimulation requires parvalbumin interneuron-dependent hippocampal neurogenesis"

**Supplementary Figures and Figure Legends**

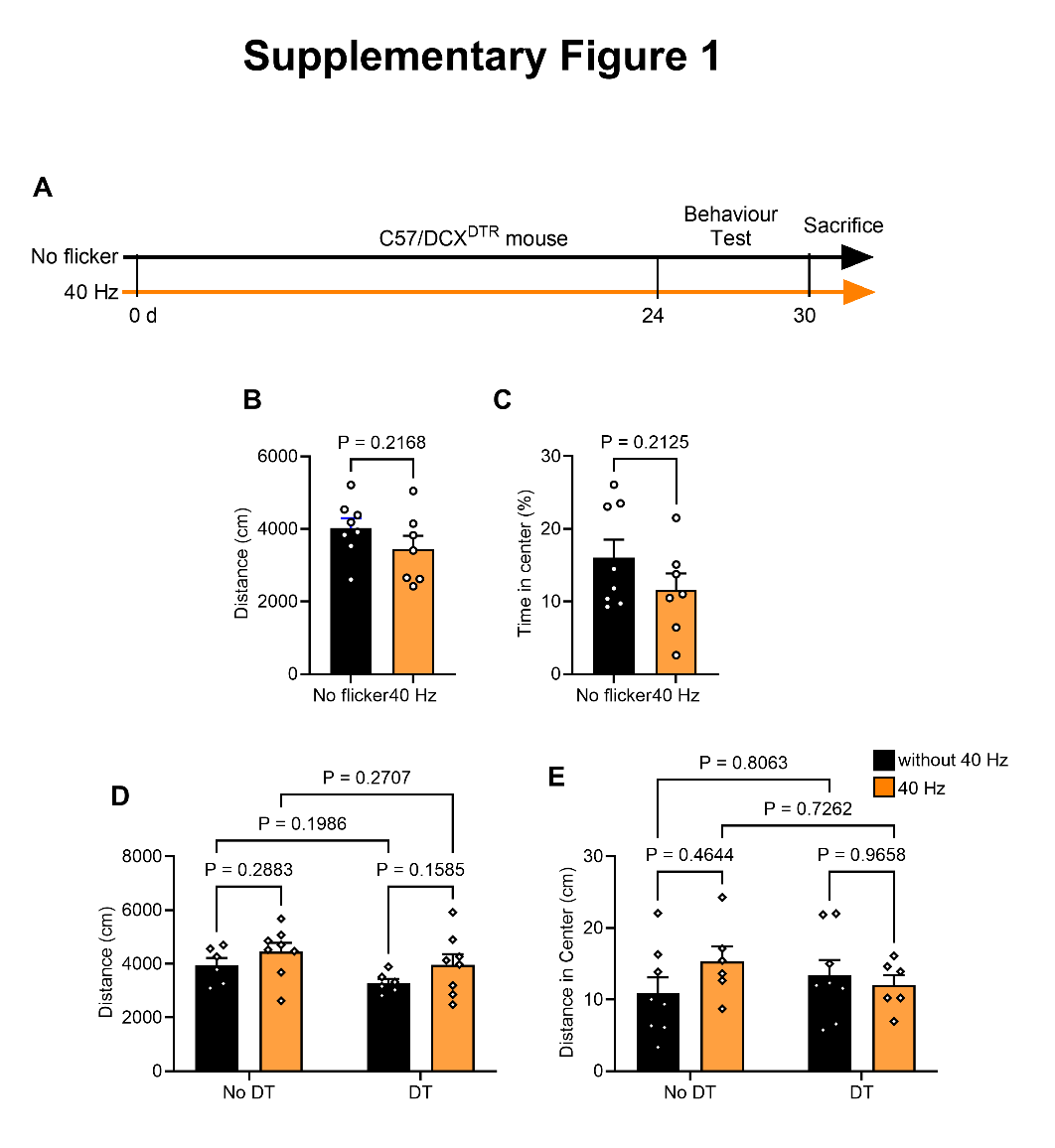

**Figure S1. Long-term 40 Hz light flicker treatment does not alter locomotion.** (A) Experimental schema for light flicker treatment (orange colored line) and APA testing. (B, C) Total distance traveled and time spent in the center were measured and compared between the No flicker and the 40 Hz light flicker-treated (40 Hz) groups. n = 8 mice in the No flicker group and n = 7 mice in the 40 Hz group. (D, E) The total distance traveled (D) and distance in the center (E) were measured and compared between the four groups of DCX^DTR^ mice. Student's t test was performed for panels B and C. A two-way ANOVA with Tukey's *post hoc* test was performed for panels D and E to show the no effect of 40 Hz light flicker treatment and the DT treatment on the distance traveled [D; F(24, 1) = 3.23, p = 0.0849] and the distance in the center [E; F(24, 1) = 0.51, p = 0.4801]. n = 8 mice in the No flicker groups and n=6 mice in the 40 Hz groups.

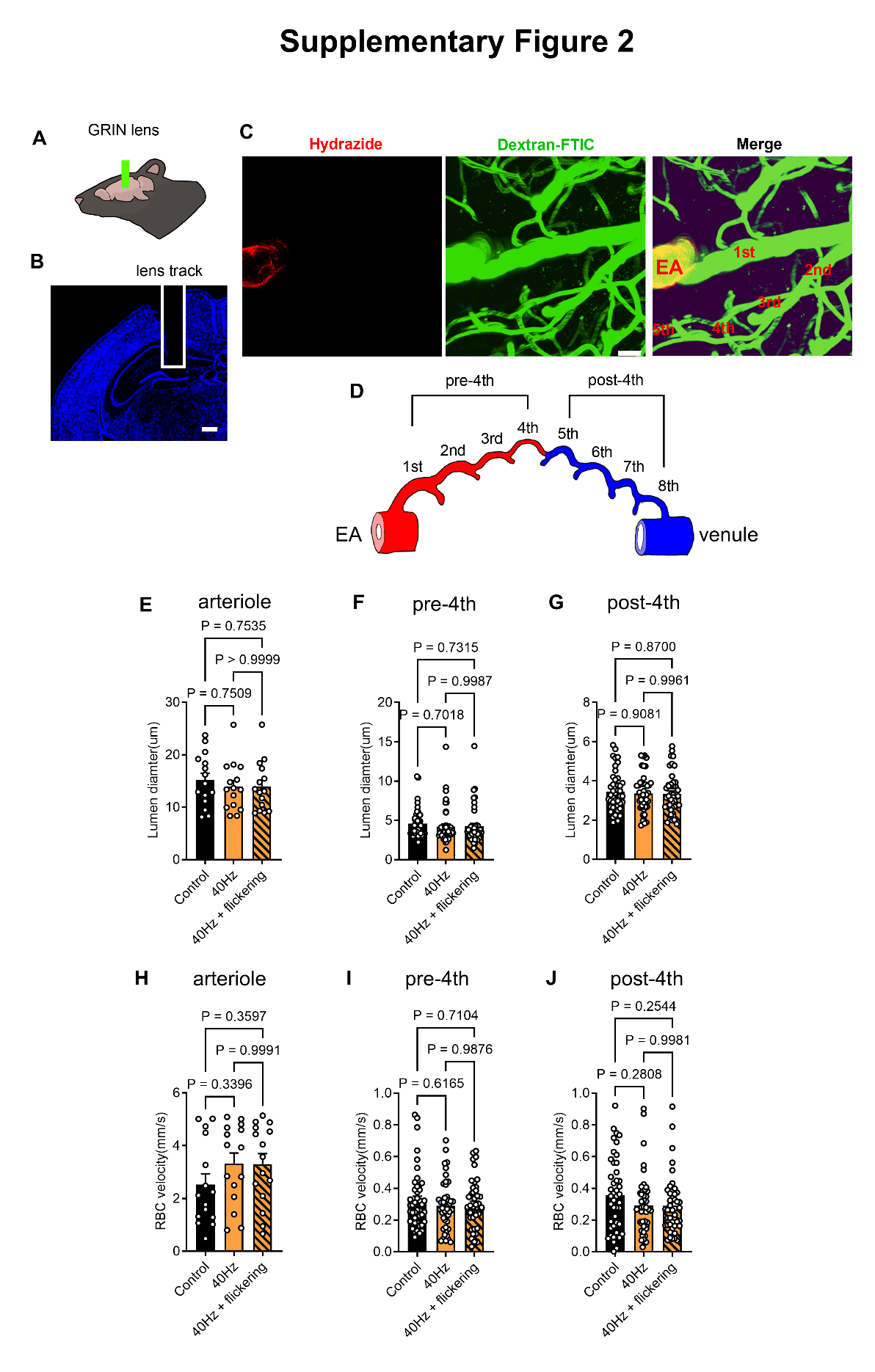

**Figure S2. Long-term 40 Hz light flicker treatment does not alter regional blood flow in the DG region.** (A) Diagram of the GRIN lens insertion in the mouse brain hippocampus. (B) The brain section indicates the exact location of the GRIN lens in the DG molecular layer (white-colored box). (C) Two-photon live imaging of the fluorescent dye Alexa Fluor™ 647 Hydrazide (red color) labeled arteriole and Dextran-FITC (green color) labeled the arterioles and the capillaries. The groups are Control (no flicker), 40 Hz (30 days of 40 Hz light flicker treatment), and 40Hz+flickering (30 days of 40 Hz light flicker treatment plus 40 Hz flickering at the time of measurement). (D) The capillary right after the ending of the hydrazide signal was defined as the 1st level capillary. Branching from the 1st level capillary produces the 2nd level capillary, and so forth. The 1st to 4th level capillaries were defined as the pre-4^th^ capillary, and the 5th to 8th as the post-4^th^ capillary. (E-G) The arteriole lumen diameter changes during 40 Hz light flicker as measured using two-photon microscopy. (H-J) The red blood cell velocity was also measured and compared as indicated. One-way ANOVA with Tukey's *post hoc* test was performed for panels from D to I. The specific P values are as indicated. Scale bar, 10 µm. n = 105 vessels from 3 mice per group.

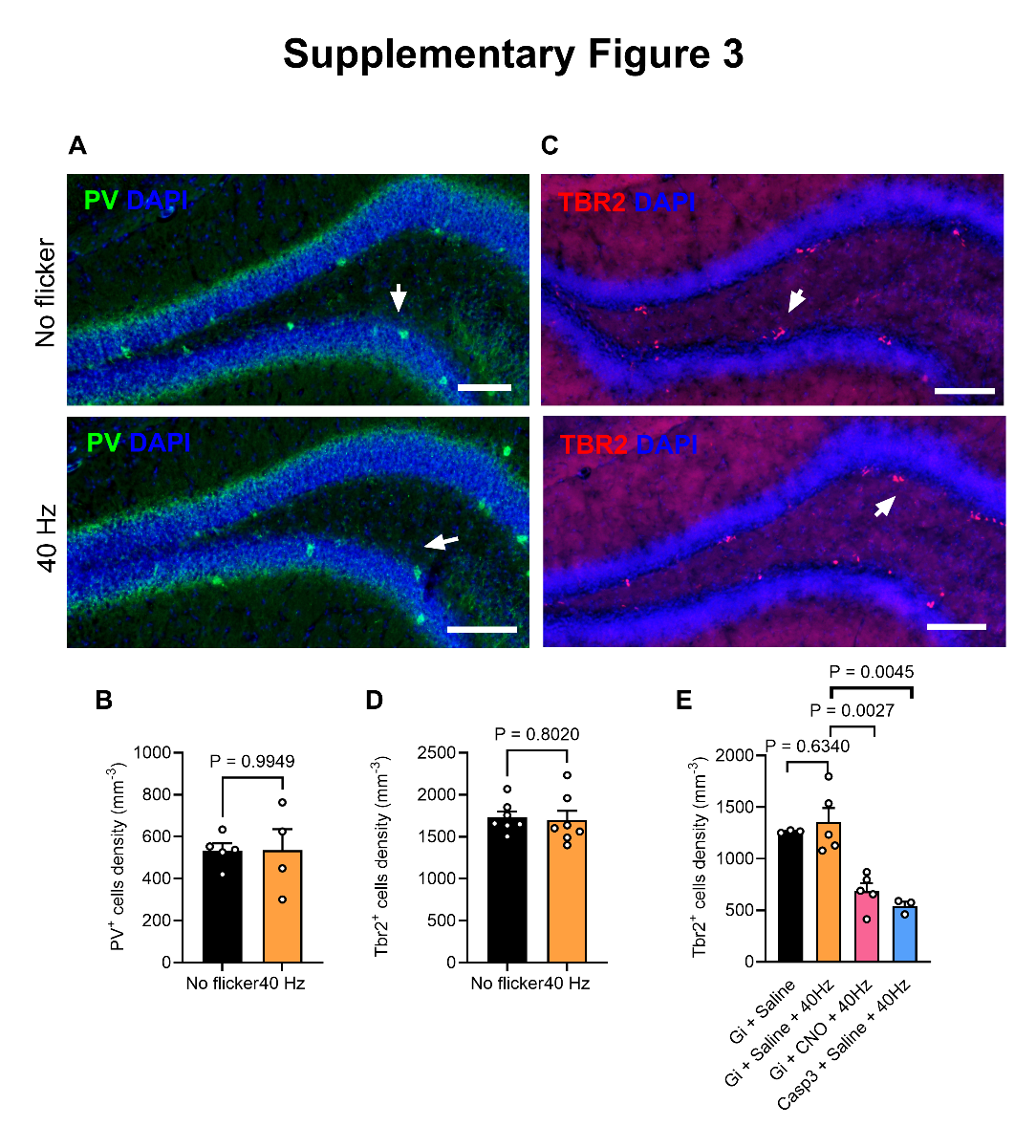

**Figure S3. The density of PV interneurons and TBR2^+^ cells in the DG remains unchanged following prolonged exposure to 40 Hz light flicker.** (A-B) Hippocampal sections PV interneurons (A, arrows) did not significantly alter the density after long-term 40 Hz light flicker treatment (B). (C-D) The density of TBR2^+^ cells (C, arrows) also did not exhibit a significant change post long-term 40 Hz light flicker treatment (D). Student's t-test was performed for panels B and D. Scale bar, 100 µm. (E) The number of TBR^+^ cells was significantly reduced in PV-inactivated (Gi+CNO+40Hz group) and PV-deleted mice (Casp3+Saline+40Hz group). The Students' t-test was performed for panel E with P values as indicated.

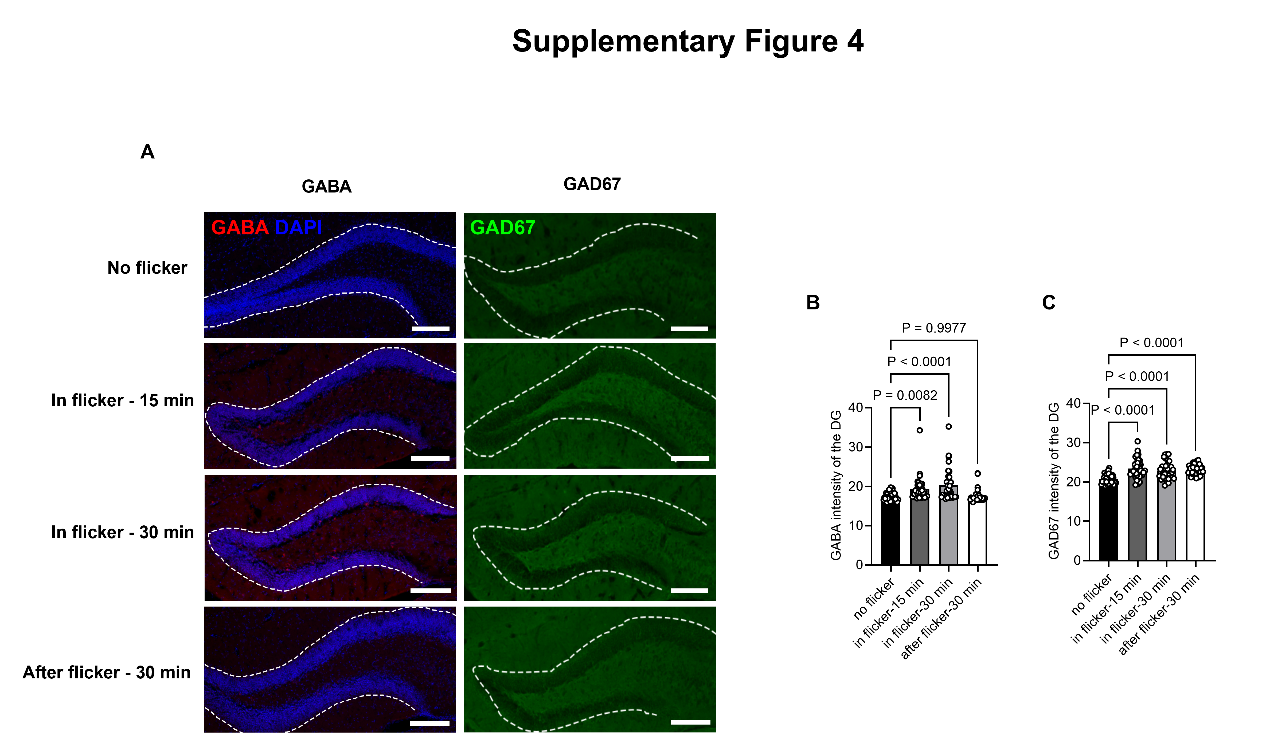

**Figure S4. Short-term 40 Hz light flicker altered GABA and GAD67 levels in the DG region.** (A) Co-staining for GABA (red), GAD67 (green), and DAPI (blue) of the DG hilus (within the white-colored dotted lines) of different time point groups. These groups were No flicker and 40 Hz light flicker treatment for 15 min (In flicker-15 min), 30 min (In flicker-30 min), and 30 min after 40 Hz light flicker treatment (After flicker-30 min). Scale bars, 100 μm; (B, C) Comparison of GABA and GAD67 intensity among different time points. One-way ANOVA with Tukey's *post hoc* test. Specific P values are as indicated on panels B and C.
